# Stability of the pH-Dependent Parallel-Stranded d(CGA) Motif

**DOI:** 10.1101/2020.07.02.184911

**Authors:** E. M. Luteran, J. D. Kahn, P. J. Paukstelis

## Abstract

Non-canonical DNA structures that retain programmability and structural predictability are increasingly being used in DNA nanotechnology applications, where they offer versatility beyond traditional Watson-Crick interactions. The d(CGA) triplet repeat motif is structurally dynamic and can transition between parallel-stranded homo-base paired duplex and anti-parallel unimolecular hairpin in a pH-dependent manner. Here, we evaluate the thermodynamic stability and nuclease sensitivity of oligonucleotides composed of the d(CGA) motif and several structurally related sequence variants. These results show that the structural transition resulting from decreasing the pH is accompanied by both a significant energetic stabilization and decreased nuclease sensitivity as unimolecular hairpin structures are converted to parallel-stranded homo-base paired duplexes. Furthermore, the stability of the parallel-stranded duplex form can be altered by changing the 5′-nucleobase of the d(CGA) triplet and the frequency and position of the altered triplets within long stretches of d(CGA) triplets. This work offers insight into the stability and versatility of the d(CGA) triplet repeat motif and provides constraints for using this pH-adaptive structural motif for creating DNA-based nanomaterials.

**STATEMENT OF SIGNIFICANCE:** This article addresses the stability of the d(CGA) triplet motif and variants in solution. Our study reveals changes in thermodynamic stability and nuclease resistance in response to pH. The identity of the 5′-nucleobase within each triplet and the position and frequency of different triplets within stretches of d(CGA) triplets can tune parallel-stranded duplex stability. This tunability can be used for nanotechnological applications where the specificity of the 5′-nucleobase pairing interaction is used to order of long stretches of d(CGA) triplets. These results can inform the rational design of pH-sensitive structurally switchable DNA-based nanomaterials.

## INTRODUCTION

Nucleobase pairing and stacking interactions inherent in self-complementary DNA provide the structural programmability and predictability needed to create reliable nanoscale self-assembling materials (1–5). Non-complementary sequences have the potential to form alternative DNA structures stabilized by non-Watson Crick base pairing and other interactions. The formation of such structures depends on local sequence and environmental factors, but the rules for stability and structure are not as established as they are for canonical duplex DNA. Predictable non-canonical structures can expand DNA’s structural versatility for nanotechnology applications (6–10). Previous work on the nanotechnological applications of non-canonical DNA have focused primarily on motifs known to be biologically relevant, including G-quadruplexes (11–13), i-motifs (14–16), and triplex-forming strands (17–19). One potential advantage of these and other noncanonical structures are their sensitivities to the local environment, including cations, salt concentration, or pH, which allow them to undergo predictable structural changes in response to environmental perturbations. (20–23).

DNA oligonucleotides containing d(CGA) triplet repeats undergo pH-dependent structural switching (Figure 1A) (24–27). Under acidic conditions, the d(CGA) motif forms a parallel-stranded duplex comprised completely of homo-base pairs (C-CH^+^, G-G, and A-A, Figure 1B) (25,26). At neutral pH, however, oligonucleotides composed of d(CGA) repeats form anti-parallel interactions that are consistent with the formation of a unimolecular DNA hairpin stabilized by G-C pairs separated by potential noncanonical A-A base pairs (24–27). At high DNA concentration and low temperature, changes in salt concentration and pH allow d(CGA)_4_ to sample four unique conformations – parallel-stranded duplex, anti-parallel hairpin, anti-parallel B-form duplex, and Z-form duplex (27). The parallel-stranded duplex is primarily observed in acidic conditions, while at higher pH (7.0 and 8.0) and low ionic strength, the structure converts to an anti-parallel hairpin. When ionic strength is increased at pH 8.0, the DNA favors an anti-parallel duplex, while very high ionic strength (4 M NaCl) with trace amounts of Ni^2+^ causes a switch in helix handedness to form Z-DNA.

**FIGURE 1.**
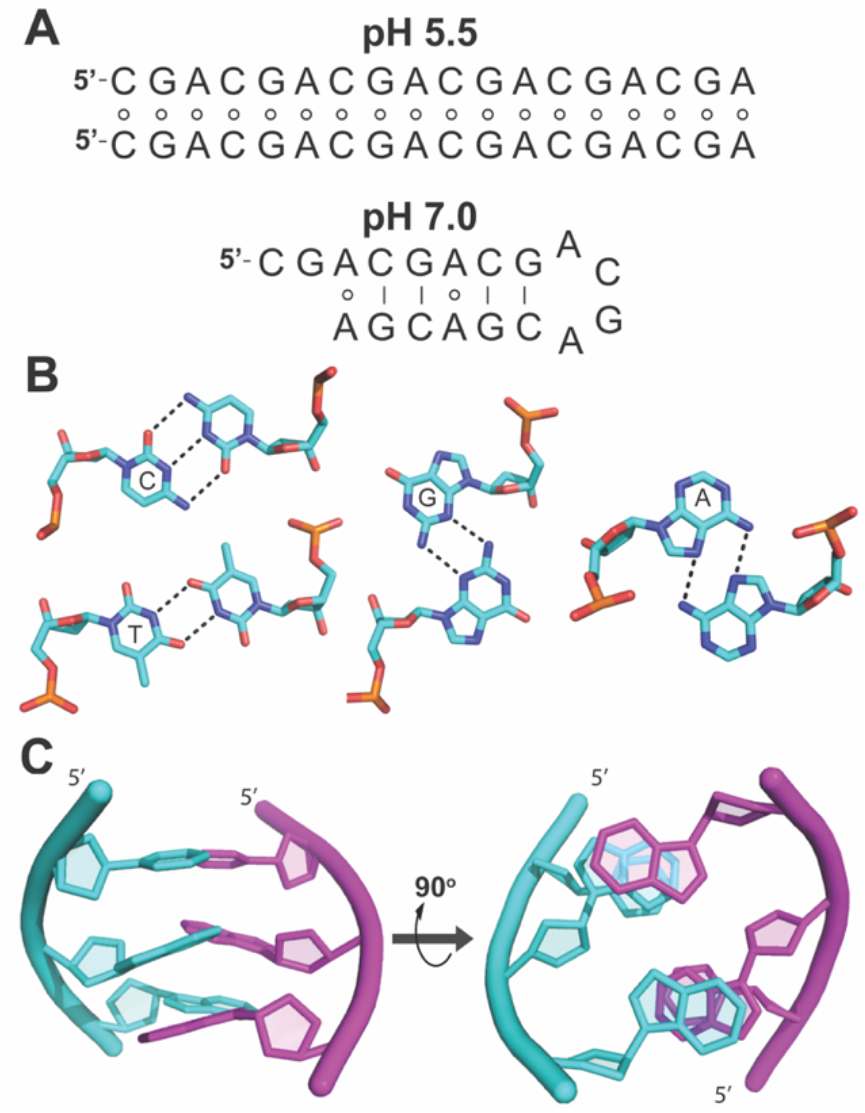
d(CGA) triplet motif. (A) (CGA)_6_ is predicted to form a parallel-stranded homo-duplex at pH 5.5 and an anti-parallel hairpin at pH 7.0, based on NMR and circular dichroism analysis (24–27). Circles indicate non-canonical base pairing and solid lines represent canonical base pairing. It is unclear whether the anti-parallel hairpin anticipated at pH 7.0 forms with (shown) or without (not shown) A-A homo-base pairs (24). (B) Homo-base pairing interactions in the parallel stranded form of the d(YGA) motif: protonation of cytosine N3 allows formation of a hemiprotonated C-CH^+^ homo-base pair; T-T homo-base pair; G-G minor groove edge homo-base pair; and A-A Hoogsteen-face homo-base pair. (C) A d(CGA) triplet forming a parallel-stranded homo-duplex, shown from the side and also rotated 90o so the 5′ ends go into the page, to show interstrand GA stacking interactions. Schematics in (B) and (C) used PDB structures 4RIM and 5CV2.

Though the d(CGA) repeat motif is capable of adopting multiple structural forms, only its parallel-stranded form has been extensively structurally characterized (23,25,28–31). Originally called II-DNA, the parallel motif is stabilized by hemi-protonated C-CH^+^, G-G minor groove edge, and A-A Hoogsteen-face homo-base pairs (Figure 1B) (26, 31). The interstrand stacking of the purine nucleobases in the GA dinucleotide steps (Figure 1C) is thought to provide a significant stabilizing force for the motif and has been observed in a number of different sequence contexts (28, 31–34). A crystal structure containing both d(CGA) and internal d(TGA) triplets revealed that interstrand GA stacking is identical in the d(TGA) triplet and the d(CGA) motif, but there is increased structural asymmetry and duplex bending associated with the C-CH^+^ base pair (29). Additionally, the same cross-strand GA dinucleotide step seen in d(CGA) and d(TGA) was observed in parallel 5′-d(GGA) sequences both in solution and in the non-canonical motif of a continuously base-paired 3D DNA lattice (33,35). In these examples the G-G base pair requires one of the two Gs to adopt a *syn* glycosidic angle to allow Watson-Crick-to-Hoogsteen face base pairing. The overall structural similarity of the d(CGA), d(TGA), and d(GGA) motifs suggests that they could be used together in d(NGA) repeat sequences, where specificity of the C-CH^+^, T-T, or G-G base pairs provide the strand registration that would be necessary for designing nanoscale DNA constructs that adopt a single base pairing geometry.

In this work, we investigate the stability of the d(CGA) motif and related sequences in solution at near-physiological temperature and salt concentration. Thermodynamics derived from UV absorbance melting curves indicate that the parallel-stranded homo-duplex structure is energetically more favorable than the anti-parallel structures, and that the frequency and identity of the 5′-nucleotide in the triplet repeat can have dramatic impacts on the relative stability. Also, increased resistance against double-strand specific nucleases in the parallel duplex form relative to the anti-parallel duplex suggests that the parallel motifs may offer unique advantages for the design, assembly, and delivery of nanostructures in cellular environments.

## MATERIALS AND METHODS

### Oligonucleotide Synthesis and Purification

DNA oligonucleotides (Table S1) were synthesized on a 1 μmol scale using standard phosphoramidite chemistry on an Expedite 8909 Nucleic Acid Synthesizer (PerSeptive Biosystems, Framingham, MA) with reagents from Glen Research (Sterling, VA). Oligonucleotides were purified by denaturing 20 % (19:1) acrylamide/bis-acrylamide, 7 M urea gel electrophoresis and dialyzed against deionized water, as previously described (35).

### Absorbance Melting Curve Procedures

UV melting curves were obtained on a Cary Bio 100 UV-visible spectrophotometer (Varian/Agilent, Palo Alto, CA) equipped with a 12-cell sample changer and Peltier heating/cooling system. DNA samples were diluted to working concentrations in either 20 mM MES pH 5.5 or 20 mM sodium cacodylate pH 7.0, both supplemented with 100 mM sodium chloride. Sample absorbance data was obtained at 260 nm as the temperature was ramped from 4 °C to 95 °C at 1 °C/min. In addition, data were collected at 260 nm for DNA renaturation from 95 °C to 4 °C at 1 °C/min to check reversibility. Self-masking quartz cuvettes with 1 cm path lengths were used for 1.4 to 20 μM samples, while a 2 mm path length cuvette was used for 50 to 120 μM samples. Final working oligonucleotide concentrations were calculated from the absorbance at 85 °C.

### Melting Curve Data Analysis

The measured absorbance values for samples at high concentration were corrected for deviations from linearity with respect to DNA concentration. A calibration curve was obtained and data falling above the linear range were corrected accordingly. We verified that this procedure yielded absorbance versus wavelength curves that were superimposable at all concentrations for samples that exhibited unimolecular behavior. Thermodynamic parameters for parallel-stranded duplex formation at pH 5.5 were obtained from the fit of each temperature versus absorbance melting curve to the bimolecular van’t Hoff expression, as described by Petersheim and Turner (36). Similarly, the van’t Hoff expression for unimolecular hairpin formation was used to fit the melt curves and extract thermodynamic parameters for data at pH 7.0 (37). The thermodynamic parameters obtained from the fits to the melting curves are reported as the average ± standard deviation of the results from independent fits from experiments with 4-6 independently prepared samples at concentrations ranging from 2-120 μM.

### Circular Dichroism (CD) Spectroscopy

CD spectra were obtained using a Jasco J-810 spectropolarimeter fitted with a thermostatted cell holder (Jasco, Easton, MD). Samples were prepared using 10 μM DNA in the same buffers used for UV melting experiments: 20 mM MES, 100 mM sodium chloride (pH 5.5), or 20 mM sodium cacodylate, 100 mM sodium chloride (pH 7.0). Samples were incubated at 4 °C overnight prior to data collection. Data were collected at room temperature at wavelengths from 220-320 nm.

### S1 Nuclease Stability Assay

20 μM DNA was incubated with 200 U/mL S1 nuclease (Thermo Fisher Scientific, Waltham, MA) in 40 mM sodium acetate pH 4.5 or 40 mM Tris HCl pH 7.5, 300 mM sodium chloride, and 2 mM zinc sulfate at 22 °C for 2 hr. The reaction was quenched with 30 mM EDTA and incubated at 70 °C for 10 min. Samples were analyzed by denaturing 20 % (19:1) acrylamide/bis-acrylamide, 7 M urea polyacrylamide gel electrophoresis and stained with SYBR Gold (Thermo Fisher Scientific, Waltham, MA).

### DNase1 Nuclease Stability Assay

DNA was incubated overnight at 4 °C with 50 mM magnesium formate to induce structure formation, as previously described (38). The DNA was diluted to 40 μM and was incubated with 7 U/mL DNase I (Thermo Fisher Scientific, Waltham, MA), 10 mM MES pH 5.5 or 10 mM Tris pH 7.5, 2.5 mM magnesium chloride, 0.5 mM calcium chloride at 37 °C for 10 min. The reaction was quenched by addition of 5 mM EDTA and incubated at 75 °C for 10 min. Samples were analyzed by 20 % denaturing polyacrylamide gel electrophoresis as above and stained with SYBR Gold (Thermo Fisher Scientific, Waltham, MA).

## RESULTS AND DISCUSSION

### (CGA)_6_ adopts parallel-stranded duplex and anti-parallel hairpin forms in response to pH

Previous studies demonstrated that d(CGA)_2_ and d(CGA)_4_ repeat sequences could transition between anti-parallel hairpin and parallel-stranded duplex (27). To determine if this behavior is also found for (CGA)_6_, we analyzed the oligonucleotide and related variants by circular dichroism and UV absorbance melting (Table 1). The CD spectrum for (CGA)_6_ at pH 5.5 showed a prominent positive band at 265 nm and a negative band at 245 nm, consistent with parallel-stranded homo-duplex formation (Figure 2A) (26,27). A clear apparently two-state UV absorbance melting transition was observed for (CGA)_6_ at pH 5.5, with a concentration-dependent melting temperature (T_m_) indicating a reaction molecularity greater than one (Figure 2B), consistent with the parallel bimolecular complexes seen in crystal and solution structures containing the d(CGA) motif (25–29). At pH 7.0, the measured T_m_ is independent of concentration, indicating a unimolecular structure, and the CD spectrum is characteristic of anti-parallel strands (Figure 2A, B). These results are consistent with (CGA)_6_ forming anti-parallel hairpin structures at neutral pH and parallel-stranded homo-duplexes under acidic conditions.

**TABLE 1.**
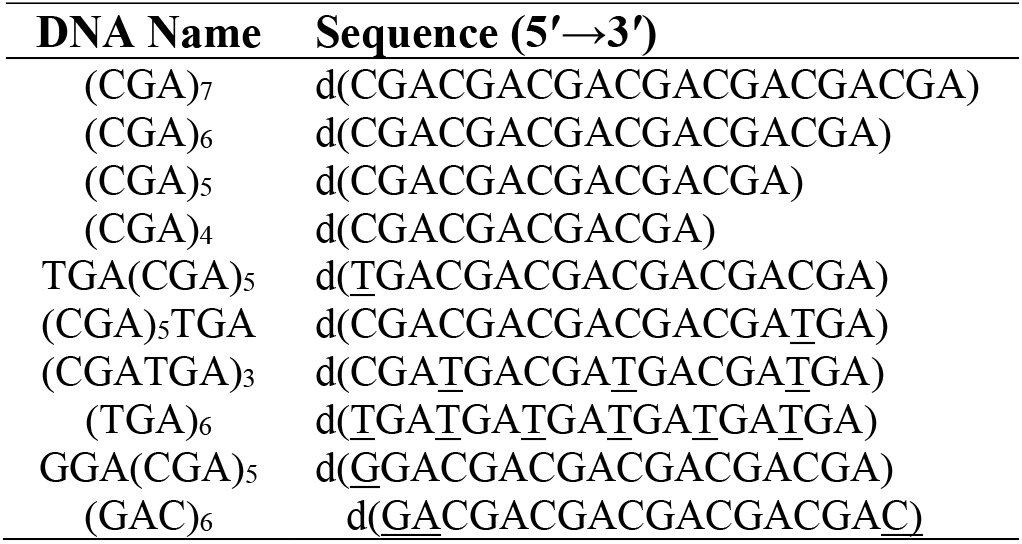
Sequences of DNA oligonucleotides. Underlined nucleotides differ from the (CGA)_n_ pattern.

**FIGURE 2.**
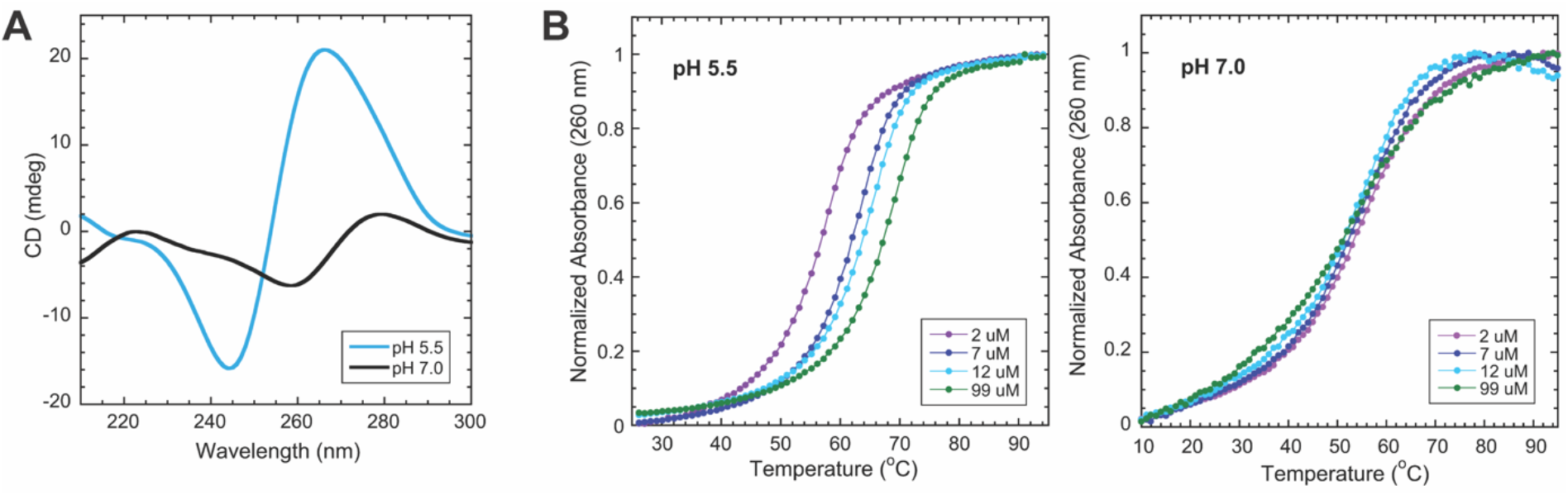
(CGA)_6_ adopts a parallel-stranded duplex or an anti-parallel hairpin in response to pH. (A) Circular dichroism spectra of 10 μM (CGA)_6_ at pH 5.5 and pH 7.0. At pH 5.5, the prominent positive band at 265 nm and negative band at 245 nm are consistent with parallel-stranded duplex formation. At pH 7.0, the positive band at 275 nm and negative band at 258 nm are characteristic of anti-parallel strands (26). (B) Normalized temperature vs absorbance curves for (CGA)_6_ show concentration dependent T_m_ at pH 5.5 and concentration independent T_m_ at pH 7.0, suggesting bi-/multi-molecular and largely unimolecular structure formation, respectively.

### Parallel-stranded duplexes are more stable than anti-parallel hairpins

(CGA)_6_ was analyzed by UV melting to establish thermodynamic parameters for the conformations observed at each pH (Table 2). Thermodynamic parameters for the parallel duplex or hairpin formation were extracted from UV melting curves fit to two-state bimolecular or unimolecular models, respectively. For each dataset, the entire absorbance vs temperature curve was fit with the van’t Hoff equation assuming constant enthalpy (ΔH°) and entropy (ΔS°), as described in Materials and Methods. While (CGA)_6_ exhibited apparent two-state melting profiles at both pH 5.5 and 7.0 and had similar melting temperatures in this concentration range, detailed examination of the pH 7.0 melting curves (Figure 2B) shows that there are significant underlying differences in the stabilities of the two structures. The higher baseline slope of the anti-parallel form at pH 7.0 suggest that its structure may be changing as a function of temperature even at low temperature. The decrease in the slope at T_m_ as concentration increases (i.e. an apparent decrease in ΔH° and ΔS°) suggests some bimolecular or multimolecular behavior. Comparing the thermodynamic parameters of (CGA)_6_ determined at pH 5.5 and at 7.0, the standard state formation free energy (ΔG°_37_) and ΔH° were destabilized by 12.2 kcal/mol and 58.6 kcal/mol, respectively; the similar T_m_s arise from the difference between unimolecular interaction at very high effective local concentration vs. bimolecular interactions at concentrations much less than the standard state 1 M. This suggests that the parallel-stranded duplex form containing intermolecular GA stacking and hydrogen bonding interactions provides substantial structural stabilization versus the anti-parallel hairpin.

**TABLE 2.** Thermodynamic parameters for (CGA)_6_ and variants at pH 5.5 and 7.0. (A) Thermodynamic parameters obtained at pH 5.5 are reported as averages of van’t Hoff analysis curve fitting results assuming bimolecular duplex formation, with oligomer concentrations between 1-120 × 10^−6^ M. Errors in ΔHo and ΔSo are reported as one standard deviation from at least 4 concentrations, and error in △G°_37_ is propagated as previously described (39–41). (B) T_m_ values are calculated for oligomer concentration of 3 × 10^−6^ M. (C) ΔΔGo37 for each sequence calculated with respect to (CGA)_5_. Negative values represent a stabilized free energy of formation, whereas positive values represent destabilized free energy of formation. (D) Thermodynamic parameters and T_m_ for pH 7.0 are reported as the average of 4 van’t Hoff curve fit datasets, assuming unimolecular hairpin formations, with oligomer concentrations between 1-10 x 10^−6^ M.

| *Thermodynamic Parameters, pH 5.5<sup>A</sup>* |  |  |  |  |  |
| --- | --- | --- | --- | --- | --- |
| Sequence | T<sub>m</sub> (°C)<sup>B</sup> | ΔG<sub>37</sub><sup>0</sup> (kcal/mol) | ΔH<sup>0</sup> (kcal/mol) | ΔS<sup>0</sup> (e.u.) | ΔΔG<sub>37</sub><sup>0</sup> (kcal/mol)<sup>C</sup> |
| (CGA)<sub>7</sub> | 61 | -15.6 ± 0.9 | -107.9 ± 7.2 | -298 ± 21 | -3.8 ± 1.0 |
| (CGA)<sub>6</sub> | 60 | -14.2 ± 0.7 | -92.1 ± 3.1 | -251 ± 9 | -2.4 ± 0.8 |
| (CGA)<sub>5</sub> | 54 | -11.8 ± 0.4 | -75.8 ± 1.4 | -207 ± 4 | 0 |
| (CGA)<sub>4</sub> | 43 | -8.9 ± 0.2 | -60.9 ± 3.5 | -168 ± 11 | +2.9 ± 0.4 |
| TGA(CGA)<sub>5</sub> | 58 | -13.4 ± 0.5 | -85.6 ± 2.4 | -233 ± 7 | -1.6 ± 0.7 |
| (CGA)<sub>5</sub>TGA | 49 | -10.3 ± 0.3 | -65.8 ± 2.2 | -179 ± 7 | +1.4 ± 0.5 |
| (CGATGA)<sub>3</sub> | 32 | -6.9 ± 0.3 | -59.7 ± 3.2 | -171 ± 10 | +4.9 ± 0.5 |
| (TGA)<sub>6</sub> | - | - | - | - | - |
| GGA(CGA)<sub>5</sub> | 56 | -13.2 ± 0.7 | -91.2 ± 2.6 | -252 ± 8 | -1.4 ± 0.8 |
| (GAC)<sub>6</sub> | 58 | -13.3 ± 0.4 | -85.8 ± 2.3 | -234 ± 7 | -1.5 ± 0.6 |
| *Thermodynamic Parameters, pH 7.0<sup>D</sup>* |  |  |  |  |  |
| Sequence | T<sub>m</sub> (°C)<sup>B</sup> | ΔG<sub>37</sub><sup>0</sup> (kcal/mol) | ΔH<sup>0</sup> (kcal/mol) | ΔS<sup>0</sup> (e.u.) | ΔΔG<sub>37</sub><sup>0</sup> (kcal/mol) |
| (CGA)<sub>6</sub> | 55 | -2.0 ± 0.2 | -33.5 ± 1.1 | -102 ± 3 | +9.8 ± 0.4 |

### (CGA)_6_ variants exhibit structural switching in response to pH

We designed several sequence variants to investigate how the triplet identity and position in hexa-repeats impact structural switching and stability (Table 1). (CGA)_6_, TGA(CGA)_5_, and GGA(CGA)_5_ were synthesized to probe the significance of the 5′-most capping nucleotides in parallel duplexes. Furthermore, sequences (CGA)_5_TGA, (CGATGA)_3_, and (TGA)_6_ were designed to test the positional and frequency compatibility of d(CGA) and d(TGA) triplets. Finally, (GAC)_6_ allowed a direct comparison with (CGA)_6_ to examine how a subtle terminal sequence permutation impacts thermodynamic stability.

CD analysis was used to examine the structural transitions of the variant sequences in response to pH. All variants except (TGA)_6_ and (CGATGA)_3_ had CD spectra consistent with the pH-dependent structures seen in (CGA)_6_ (Figure S1). Interestingly, CD spectra for (TGA)_6_ had characteristics of anti-parallel strands at both pH 5.5 and 7.0, while the spectra for (CGATGA)_3_ could represent an equilibrium between both forms. This suggests a decreased pH sensitivity and a preference for anti-parallel strands with increasing d(TGA) triplet content. All variants except (TGA)_6_ had concentration-dependent, two-state melting curves at pH 5.5 and concentration-independent melting transitions at pH 7.0, indicating bimolecular and intramolecular interactions, respectively. Interestingly, (CGA)_6_ was the only oligomer that exhibited clear two-state melting at pH 7.0, with all of the other sequence variants having an apparent minor lower-temperature melting transition between 20-40 °C. The multi-state transitions at pH 7.0 may reflect populations of hairpin and anti-parallel duplex structures (24).

### Anti-parallel homogeneity correlates with d(CGA) repeat length

At pH 7.0, there are interesting differences in the melting profiles for (CGA)_n_ (n = 4 – 7) (Figure S2,S3). When n = 4 or 5, the melting profile is broad with multiple transitions, whereas when n ≥ 6 the melting transition is still broad but becomes more two-state. The multi-transition melting profile seen in shorter (CGA)_n_ repeats (n ≤ 5) suggests that a temperature dependent equilibrium between hairpin and duplex or higher-order forms may exist. In contrast, the two-state profile for longer repeat sequences (n ≥ 6) suggest they exist predominantly in the hairpin conformation. This trend is similar to the length-dependent conformational equilibrium seen in other triplet repeats (24) and further supports the importance of oligo lengths in rational structure design.

### Estimated ΔG°_37_ values can be used to predict the energetics of new triplet repeat sequences

To estimate the energetic contribution of each parallel-stranded triplet unit we compared thermodynamic parameters of oligonucleotides with differing number of repeat units. Strong apparent enthalpy-entropy compensation is observed for the set of (CGA)_n_ variants tested (Figure S4). The positive correlation seen in the ΔH° vs ΔS° plot demonstrates the strong enthalpy-entropy compensation, while the ΔG°_37_ vs ΔH° plot confirms that the apparent compensation is not an artifact of experimental error. As a simple predictor of the ΔG°_37_ for (CGA)_n_ sequences, the number of d(CGA) units were plotted against the respective experimental ΔG°_37_ value (Figure S5). Interestingly, the *y*-intercept (−0.250 kcal/mol) of the linear fit, which corresponds to the ΔG°_37_ for a n=1 sequence, could represent the gain in stability upon formation of the first triplet unit slightly overbalancing the entropic penalty of bringing the strands together. The linear relationship can be used to predict the ΔG°_37_ for a (CGA)_n_ triplet of any length and can be combined with other estimates (5′-d(GGA), 5′-d(TGA), 3′-d(TGA), see below) to predict ΔG°_37_ of various sequences (Figure S5). Importantly, these results are internally consistent and provide a baseline for understanding the energetic contributions of these triplets.

### The 5′-nucleotide influences parallel-duplex stability

We compared ΔG°_37_ for variants with different 5′-terminal capping triplets (d(CGA), d(GGA), or d(TGA)) to (CGA)_5_ to understand how the identity of the 5′ triplet contributed to thermodynamic stability of the parallel-stranded duplex. The 5′-d(GGA) triplet (ΔΔG°_37_ = −1.4 kcal/mol with respect to (CGA)_5_) and the 5′-d(TGA) triplet (−1.6 kcal/mol with respect to (CGA)_5_) show similar contributions, both being less stabilizing than the 5′-d(CGA) triplet, estimated above at −2.4 kcal/mol. If the 5′ triplet contains the anticipated GA interstrand stacking interactions, the differences in ΔG°_37_ between d(CGA), d(GGA), and d(TGA) triplets can be attributed primarily to the terminal 5′-nucleotide identity. Though the 5′-d(GGA) triplet appears compatible with 5′ d(CGA) in the parallel-stranded duplex, incurring only a slight thermodynamic destabilization, our preliminary modeling suggests that stretches of internal d(GGA) triplets would not be well accommodated due to structural clashes that would arise from an internal G_*syn*_ residue (33,35). CD data further suggest that stretches of d(GGA) triplets do not form the expected structure at either pH (Figure S6). The small change between the terminal d(GGA) and d(TGA) suggests that the 5′-G and 5′-T are thermodynamically comparable in the parallel duplex. Interestingly, the entropy of formation for GGA(CGA)_5_ and (CGA)_6_ is considerably lower than for TGA(CGA)_5_, which could result from conformational variability introduced by a G residue in the *syn* conformation, as seen in crystal structures containing the parallel-stranded d(GGA) motif. The destabilization caused by the 5′-T can also be attributed to enthalpic destabilization (−92.1 to −85.6 kcal/mol) reflecting the loss of one hydrogen bond by replacing the hemi-protonated C-CH^+^ with T-T base pair. Additional electronic effects from the cationic C-CH^+^ base pair that are not found in the T-T or G-G base pairs may also contribute.

Notably, the △G°_37_ for (GAC)_6_ (−13.3 kcal/mol) is different than (CGA)_6_ (−14.2 kcal/mol), though d(GAC) is a simple permutation of d(CGA). The structural basis for this difference is not immediately clear, but it does suggest that the identity of the terminal homo base pairs – 5′ or 3′ – can influence parallel stranded duplex stability. The variation in thermodynamic stability suggests that the strength of interactions at the 5′-terminus could play an important role in parallel-stranded duplex nucleation and stabilization. Nucleotides providing strong intermolecular interactions could help to lock the duplex in place to avoid premature end fraying, contributing to enhanced overall thermodynamic stability. Further studies will show whether the apparent non-nearest neighbor effects in d(GAC) vs d(CGA) act at the triplet level or the oligonucleotide terminus level.

### d(TGA) triplet position and frequency impacts the thermodynamic stability of mixed (YGA)_6_ oligomers

The identity of the 5′-pyrimidine in the d(CGA) triplet motif leads to subtle but significant structural changes in the parallel-stranded duplex, evident in crystal structures containing d(CGA) and d(TGA) repeats (29,30). C-CH^+^ base pairs induce an asymmetry, likely as a result of a hydrogen bond between N4 of one cytosine to the non-bridging phosphate oxygen of the previous intrachain adenosine (29). In contrast, the T-T base pair retains duplex symmetry at the expense of stacking interactions. Our results show that the frequency of d(TGA) triplets within (YGA)_6_ oligomers has a significant impact on thermodynamic stability of the parallel-stranded duplex.

At pH 5.5, (CGA)_6_ has a clear two-state melting transition, indicating structure formation when the sequence is completely comprised of d(CGA) triplets (Figure 3). In contrast, (TGA)_6_ had no evident melting transition at pH 5.5 or 7.0 (Figure 3), suggesting that the d(TGA) triplet is unable to adopt a stable structure at either pH. This could indicate that the interactions formed by the T-T base pair are either not strong enough to nucleate structure formation or is transient.

**FIGURE 3.**
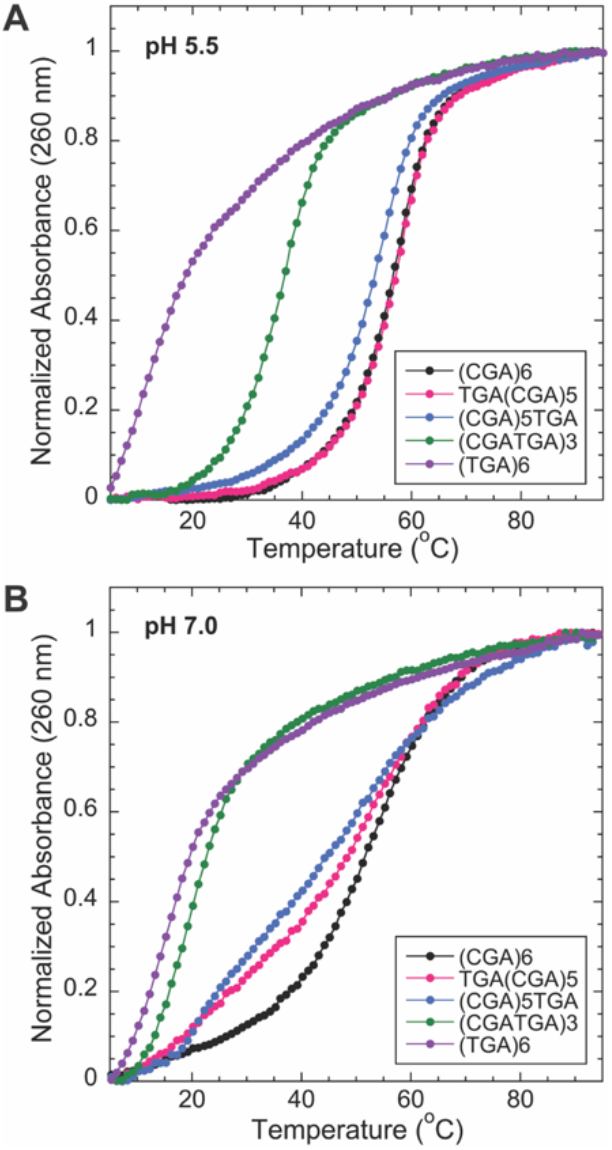
The addition of d(TGA) triplets decreases melting temperature of (YGA)_6_ oligomers. Normalized temperature vs absorbance curves for (YGA)_6_ oligomers with d(TGA) triplets (n = 6, 3, 1, 0). (A) At pH 5.5, all variants except (TGA)_6_ exhibit two-state melting. (B) At pH 7.0, (CGA)_6_ is the only variant to retain two-state melting. (CGA)_5_TGA and TGA(CGA)_5_ have broad melting curves with multiple transitions, while (TGA)_6_ and (CGATGA)_3_ do not have clear melting transitions, suggesting a lack of stable structure formation.

We tested the destabilizing effects associated with position and number of d(TGA) triplets in a given (YGA)_6_ sequence using the sequence variants TGA(CGA)_5_, (CGA)_5_TGA, (CGATGA)_3_, and (TGA)_6_. As described above, in the parallel duplex form, the addition of a d(TGA) triplet to the 5′-end stabilized the free energy of formation by −1.6 ± 0.7 kcal/mol, compared to (CGA)_5_ (Table 1). (CGA)_5_TGA was tested to observe the impact of the d(TGA) triplet at the 3′-end. This resulted in a destabilization of the free energy of duplex formation by 1.4 ± 0.7 kcal/mol, compared to (CGA)_5_, suggesting that the position of the d(TGA) triplet significantly impacts the thermodynamic stability of the parallel-stranded duplex. When the d(TGA) is at the 5′-end, the T-T homo-base pair interaction induces a slight thermodynamic destabilization, compared to a 5′-C-CH^+^ homo-base pair. In contrast, when the T-T homo-base is pair positioned internally within the 3′ triplet, there is a significantly greater impact on duplex stability, presumably through disruption of the downstream G-G and A-A base pairs. Correspondingly, more substantial destabilization (7.5 kcal/mol) was observed when multiple d(TGA) triplets were added as in (CGATGA)_3_.

Although the d(TGA) triplets decrease thermodynamic stability, they have the potential to modulate pH sensitivity. CD results for (TGA)_6_ (Fig. S1) suggest that the presence of multiple d(TGA) triplets correspond to decreased pH sensitivity and preference for the anti-parallel form. d(TGA) triplets can be combined with other strong parallel-strand inducting triplets, such as d(GGA) or d(CGA), potentially providing a unique system for tuning structure formation over specific pH ranges. Information on the thermodynamic stability of sequences containing tandem d(YGA) triplets can be combined with structural data to provide valuable insight in optimizing parallel-stranded duplexes for 3D DNA crystal design and nanoscale applications.

### Parallel-stranded duplexes resist nuclease digestion

With increasing interest in using DNA nanotechnology in cellular applications, we assessed the stability of DNA structures in conditions that mimic cellular environments. To gain insight on the behavior of the d(YGA) triplet motif in cellular conditions, we qualitatively examined resistance to nucleases with preferences for double stranded B-DNA (DNase I) or unpaired, single-stranded DNA (S1 nuclease). Control experiments confirmed the activity of both enzymes for different substrates at the pHs tested: DNase I functioned optimally on double-stranded substrates at low pH, while S1 nuclease showed decreased activity at pH 7.5 for its single-stranded substrate (Figure S7).

Each variant tested was incubated with DNase I at pH 5.5 and 7.5 (Figure 4A). At pH 7.5, (CGA)_6_, TGA(CGA)_5_, and (CGA)_5_TGA show clear cleavage patterns, suggesting that the structure formed at pH 7.5 has DNase I-recognizable regions. This supports previous analysis indicating the formation of an anti-parallel hairpin at near-neutral pH. Importantly, these same variants showed clear DNase I resistance when the reaction was repeated at pH 5.5. This suggests that the parallel-stranded duplex formed by (CGA)_6_, TGA(CGA)_5_, (CGA)_5_TGA, and (CGATGA)_3_ is not recognized by DNase I. (CGATGA)_3_ appears to have slight protection at the lower pH, while the cleavage of (TGA)_6_ is identical at both pHs. Overall, these results are consistent with our UV melting observations showing no stable structure formation for either of these last two oligos at pH 7.0 and some weakly forming structure for (CGATGA)_3_ at the lower pH.

**FIGURE 4.**
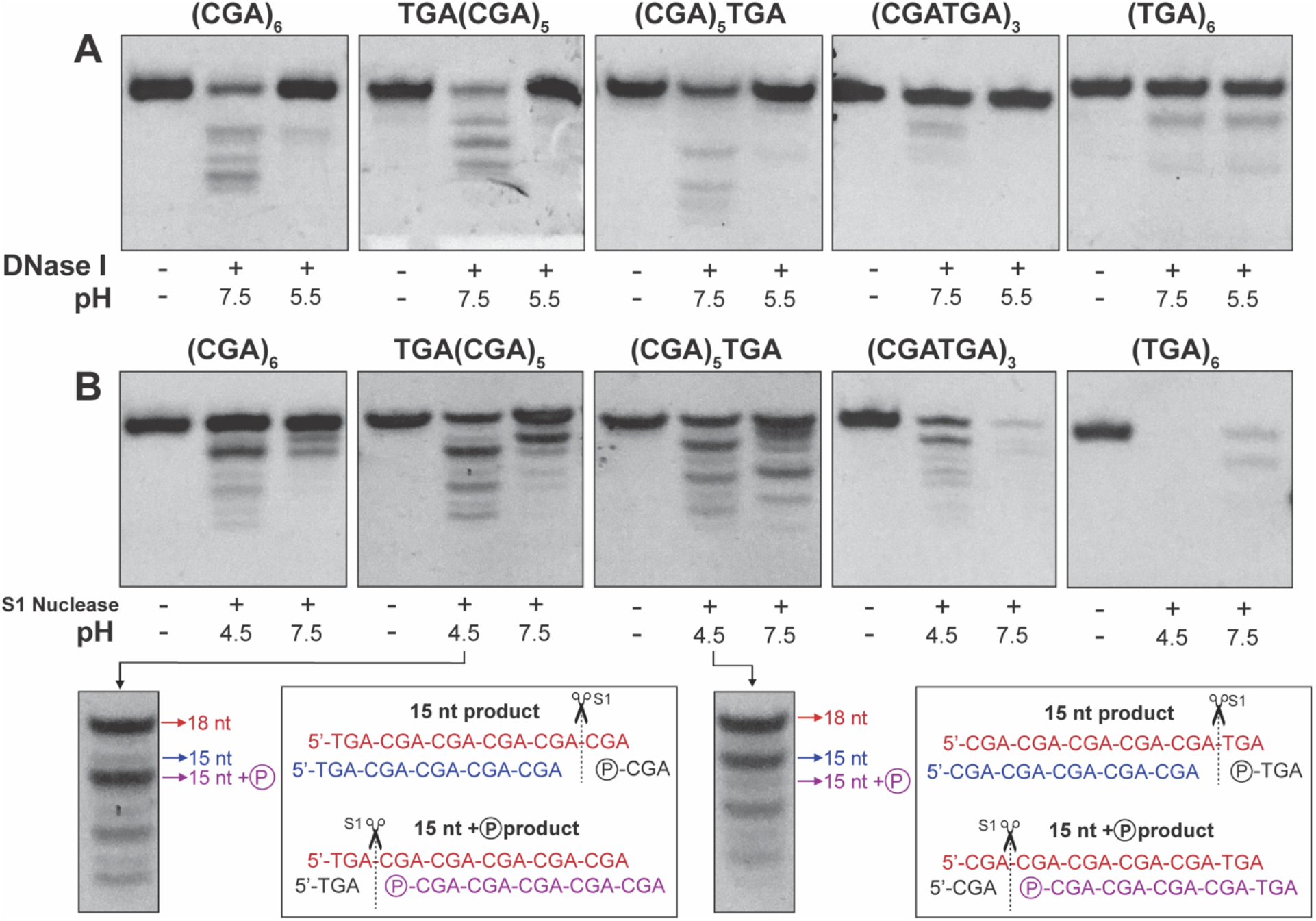
Parallel-stranded duplex enhances nuclease resistance. (A) (YGA)_6_ variants digested with DNase I at pH 7.5 or 5.5. DNase I preferentially cuts double stranded DNA, suggesting that the parallel duplexes formed at pH 5.5 are resistant to DNase I degradation. (B) (YGA)_6_ variants digested with S1 nuclease at pH 7.5 or 4.5. Red labeled sequences correspond to full length DNA (18-nt), without any nuclease modification. Blue sequences represent the nuclease cleavage product without a terminal phosphate group. The magenta sequence corresponds to the nuclease cleavage product containing a terminal phosphate group, leading to increased mobility.

A similar experiment was performed to test d(YGA) repeat susceptibility to the single-strand specific S1 nuclease. (CGA)_6_, TGA(CGA)_5_, (CGA)_5_TGA, and (CGATGA)_3_ S1 nuclease cleavage products at pH 7.5 suggest single-stranded regions are primarily associated with hairpin ends (Figure 4B). Intense bands visible at ≥14 nt are consistent with the degradation of terminal non-base paired nucleotides and or single-stranded triplet overhangs at the hairpin terminus. The difference in degradation pattern between all variants suggests that the hairpins formed are not structurally identical. All oligonucleotides were fully digested at pH 7.5 when exposed to S1 nuclease for 2 days, suggesting that the anti-parallel hairpin is dynamic enough to be susceptible to S1 digestion over extended time periods (Figure S8).

(CGA)_6_, TGA(CGA)_5_, (CGA)_5_TGA, and (CGATGA)_3_ were only partially digested by S1 nuclease at its optimal pH of 4.5 (Figure 4B). UV melting and CD experiments indicate that these oligonucleotides form parallel stranded duplexes in conditions similar to S1 nuclease reaction conditions, explaining the lack of complete S1 digestion. The intense 15-nt band present in the degradation of (CGA)_6_ suggests that a small population of the parallel-stranded duplexes are frame-shifted by one d(CGA) triplet, creating 3-nt S1 nuclease-accessible single-stranded overhangs. The intense unreacted band remaining at 18-nt suggests that most oligomers in the reaction populated the perfectly aligned parallel-stranded duplex which resisted S1 nuclease digestion. The low molecular weight degradation products could result from frame-shifted duplexes containing more than one triplet overhang. Apparently, once duplexes are formed frame shifting is extremely slow at room temperature, else equilibration among frames would have exposed all of the sequence to S1 activity.

TGA(CGA)_5_ and (CGA)_5_TGA show S1 degradation patterns similar to each other, consistent with triplet cleavage, as intense bands persist at 15, 12, and 9-nt. Interestingly, each intense band is coupled with a faint band, where the position and intensity can be used to gain insight into the S1 cleavage site. The cleavage of a 5′-triplet will produce a fast migrating 15-nt product containing a terminal phosphate, whereas, the cleavage of a 3′-triplet will produce a slower migrating 15-nt product that does not have a 5′-phosphate. The faster migrating 15-nt band is very intense in TGA(CGA)_5_, suggesting that the 5′ d(TGA) triplet was preferentially digested. The opposite is seen in (CGA)_5_TGA, where the intense 15-nt band migrates slower, suggesting that the 3′-d(TGA) was cleaved (Figure 4B). The bands at 18, 15, and 12-nt persist when exposed to S1 nuclease for up to 2 days, indicating that the parallel-stranded duplex is stable at room temperature (Figure S8). (TGA)_6_ was almost completely digested by S1 nuclease by the end of the reaction time, indicating the lack of structure formation at each pH (Figure 4B). This further confirms results from CD and UV melting experiments that suggest (TGA)_6_ does not form stable structure at acidic or neutral pH.

### Stable high molecular weight structures form in all cytosine containing variants

Unexpected high molecular weight (HMW) products form over time for (CGA)_6_ and all variants, except (TGA)_6_, and persist in 8 M urea denaturing gels (Figure S9). The formation of the stable HMW structures likely rely on cytosine reactivity, as HMW bands are never observed for (TGA)_6_ or (TGA)_5_. We cannot confirm the identity or mechanism of HMW band formation, but we suspect that stable aggregates or crosslinks are forming from degradation products (depurination) of the d(CGA) triplet repeats over time.

## CONCLUSION

This work uses a systematic approach to determine the stability of the d(CGA) triplet repeat motif and variants in solution. It is evident that (CGA)_6_ and variants undergo pH-induced structural switching that coincides with a thermodynamic destabilization. This trend can be linked to the loss of strong interstrand GA base stacking and homo-base pair interactions as pH is increased. Previous studies have investigated the formation of the parallel-stranded duplex of short (CGA)_n_ repeats (n ≤ 4) in solution (27) or within a complex crystal structure (29,31). Additional structural studies on sequences containing (CGA)_n_ triplets (n > 4) will be useful in providing in-depth insight into the behavior of long stretches of tandem d(CGA) repeats to further confirm the specific interactions responsible for the difference in thermodynamic stability at pH 5.5 and 7.0. It is clear that the addition of d(TGA) repeats decreases thermodynamic stability, but structure is not completely destroyed if d(TGA) units are in tandem with strong parallel-strand inducing triplets. (CGA)_6_ and related variants have been proposed to be useful in designing new parallel-stranded DNA architectures. Specifically, particles decorated with oligonucleotides containing d(CGA)_n_ triplets could be localized or separated based on structural changes caused by pH (Figure 5). Considering slight structural and thermodynamic deviations between each terminal 5′-residue, this motif could also serve as a discriminator in programmable pairing of long stretches of d(YGA) triplets. Additionally, the parallel-stranded duplex formed in acidic pH offers increased nuclease resistance, furthering the potential for this material to be used in cellular applications. The solution stability data presented in this work can be used to strategically design and optimize sequences for use in the rational design of 3D DNA crystals or nanotechnology applications that could benefit from pH-triggered structural switching.

**FIGURE 5.**
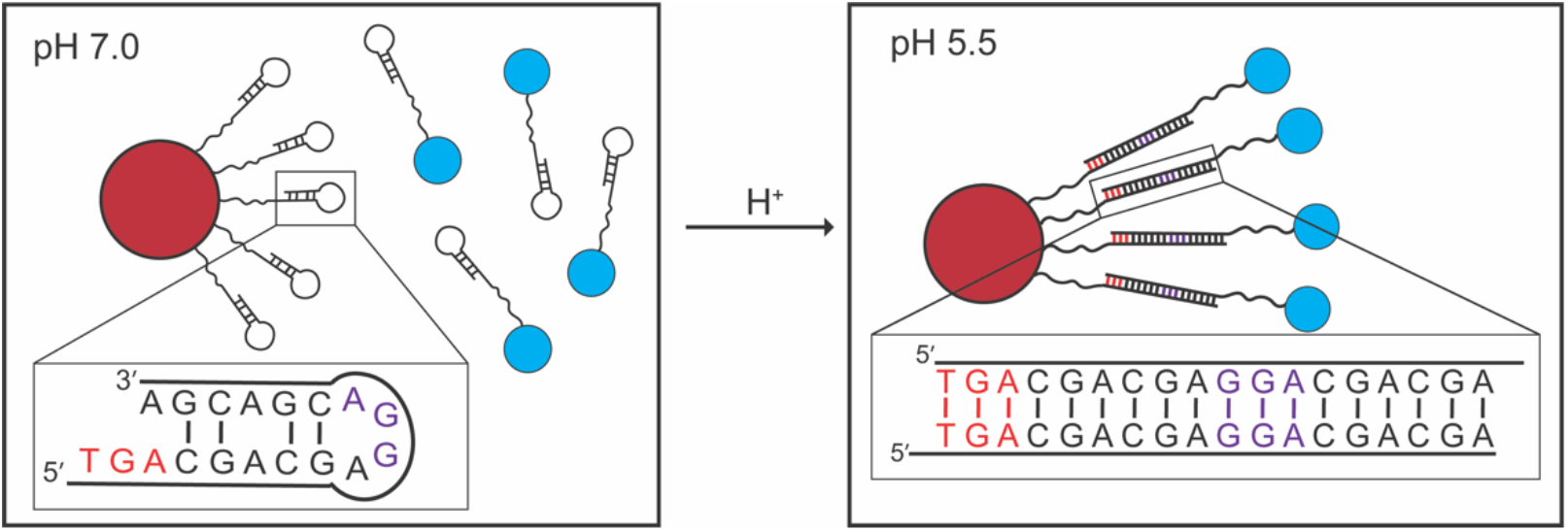
Potential use of (CGA)_6_ and variants in DNA nanotechnology applications where pH changes could localize particles together. At pH 7.0, d(CGA) forms hairpin structures, preventing duplex formation and particles remain separate. At pH 5.5, parallel-stranded duplexes can form and localize particles of interest together. d(TGA) and d(GGA) triplets can be used to ensure desired registration and length of the linking duplex. Linker stretches solely comprised of d(CGA) triplets could be used, but distance between particles of interest could vary.

## AUTHOR CONTRIBUTIONS

E.M.L. designed and performed experiments, analyzed the data, and wrote the manuscript. J.D.K. contributed to data analysis, discussion, and writing. P.J.P. designed experiments, contributed to data analysis and discussion, and wrote the manuscript.

## Supplemental Information

**FIGURE S1.**
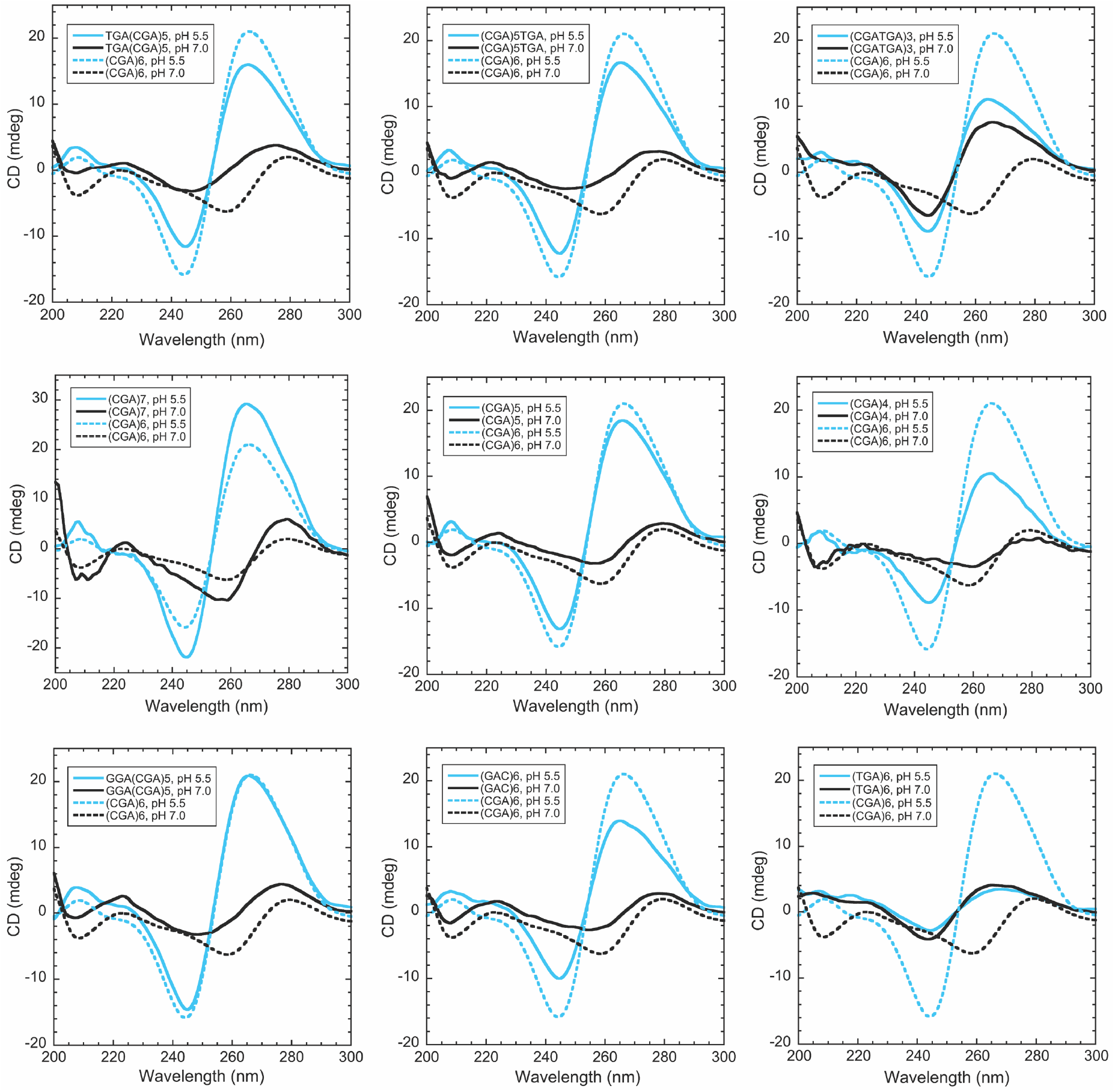
Circular dichroism spectra of 10 μM (CGA)_6_ variants. The spectra of (CGA)_6_ at pH 5.5 and pH 7.0 are shown (dashed lines) in each panel for reference. At pH 5.5 (blue curves), the prominent positive band at ~265 nm and the negative band at ~245 nm are characteristic of parallel-stranded duplex formation. As the number of d(CGA) triplets decreases, the intensity of the bands associated with parallel-stranded duplexes also decreases. At pH 7.0 (black curves), the positive band at ~275 nm and negative band at ~258 nm are characteristic of anti-parallel strands. (TGA)_6_ and (CGATGA)_3_ have bands characteristic of anti-parallel strands at each pH, suggesting that pH sensitivity is decreased or eliminated with increasing d(TGA) content.

**FIGURE S2.**
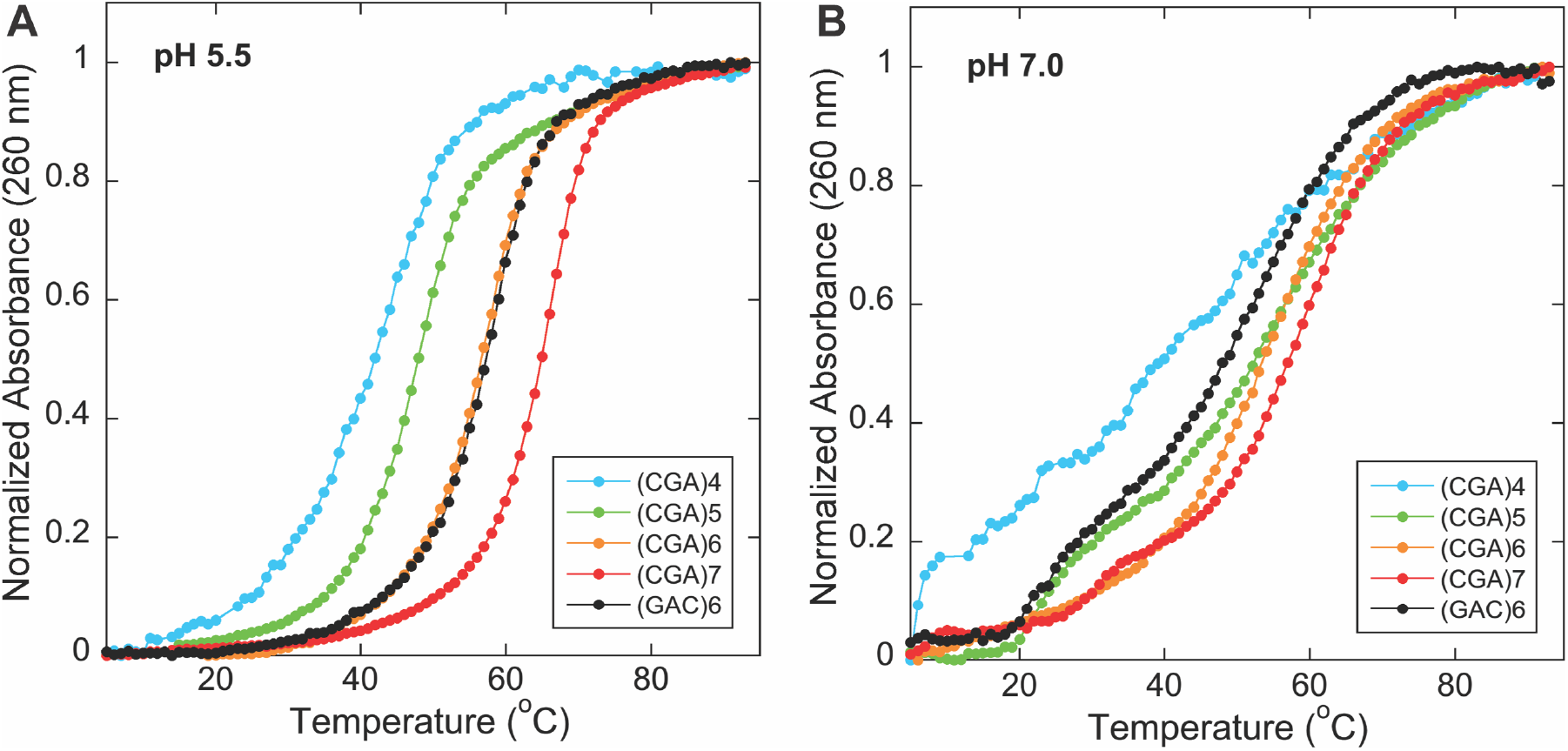
Normalized temperature versus absorbance curves for 2.5 μM (CGA)_n_ variants, where n = 4 to 7 at (A) pH 5.5 and (B) pH 7.0. It is unclear why (GAC)_6_ behaves differently than (CGA)_6_, as it is a permutation of d(CGA), at pH 7.0. We suspect that the difference could arise from variations in hairpin loop and stem size, as well as the likelihood of interconversion between different hairpin forms. Assuming the d(CGACG) loop is the most stable hairpin form, (CGA)_6_ could form a 6-bp stem where the d(CGACG) loop is capped by A-A producing the two-state transition seen at pH 7.0. The slight destabilization seen for (GAC)_6_ could arise from a similar hairpin with a d(CGACG) loop but only accommodating 5-bp. Further, the non-two state melt suggests that (GAC)_6_ could be dynamically interconverting between two different forms – one with a 5-bp stem and a d(CGACG) loop or 7-bp stem and a d(ACGA) loop.

**FIGURE S3.**
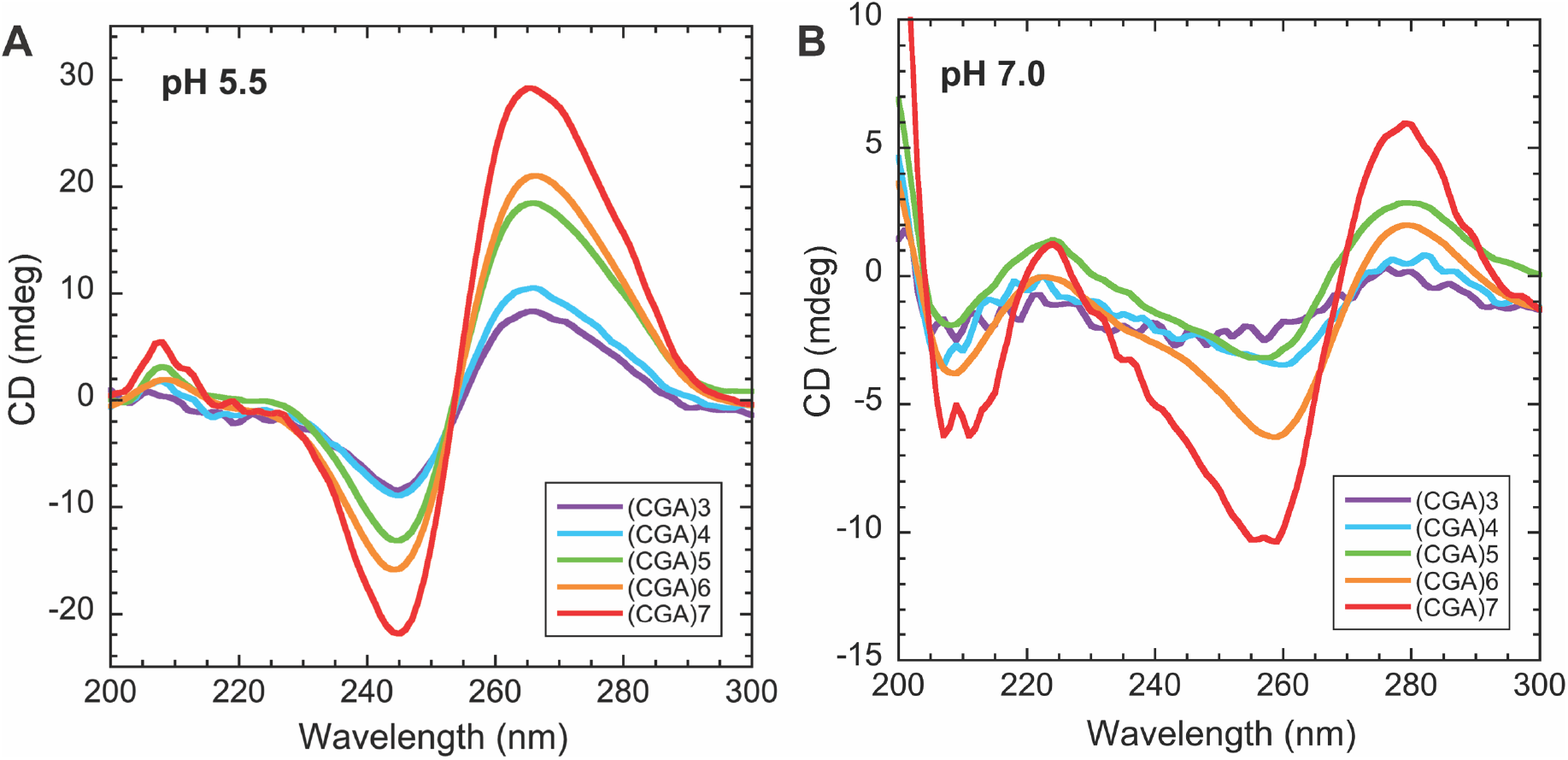
CD spectra of 10 μM (CGA)_n_ where n = 3 to 7 at (A) pH 5.5 and (B) pH 7.0. At pH 5.5, the intensity of the bands characteristic of the parallel-stranded duplex (245 nm, 265 nm) decrease as the number of d(CGA) units decrease. This suggests that the parallel-stranded form can exist at all repeat sizes tested. At pH 7.0, the signal intensity corresponding to anti-parallel strands (258 nm, 275 nm) decreases with repeat number, but is lost when n ≤ 4 suggesting that the hairpin structure cannot form at low repeat number.

**FIGURE S4.**
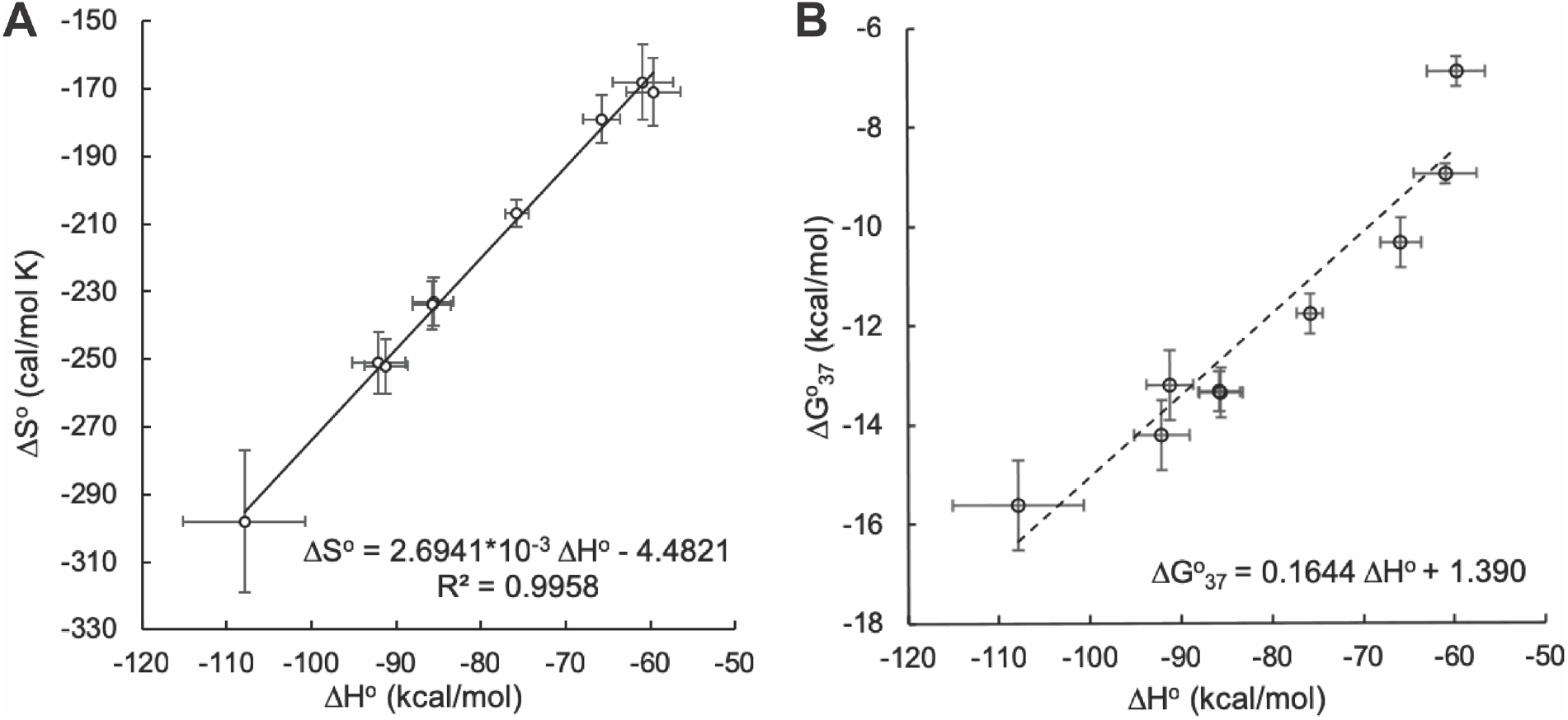
Apparent entropy-enthalpy compensation of all (CGA)_n_ variants used in this study. (A) The positive correlation between ΔH° and ΔS° indicates a strong enthalpy-entropy compensation for all oligonucleotides tested. (B) The ΔG°_37_ vs ΔH° plot suggests that compensation is due to the underlying physical reality, as opposed to experimental error.

**FIGURE S5.**
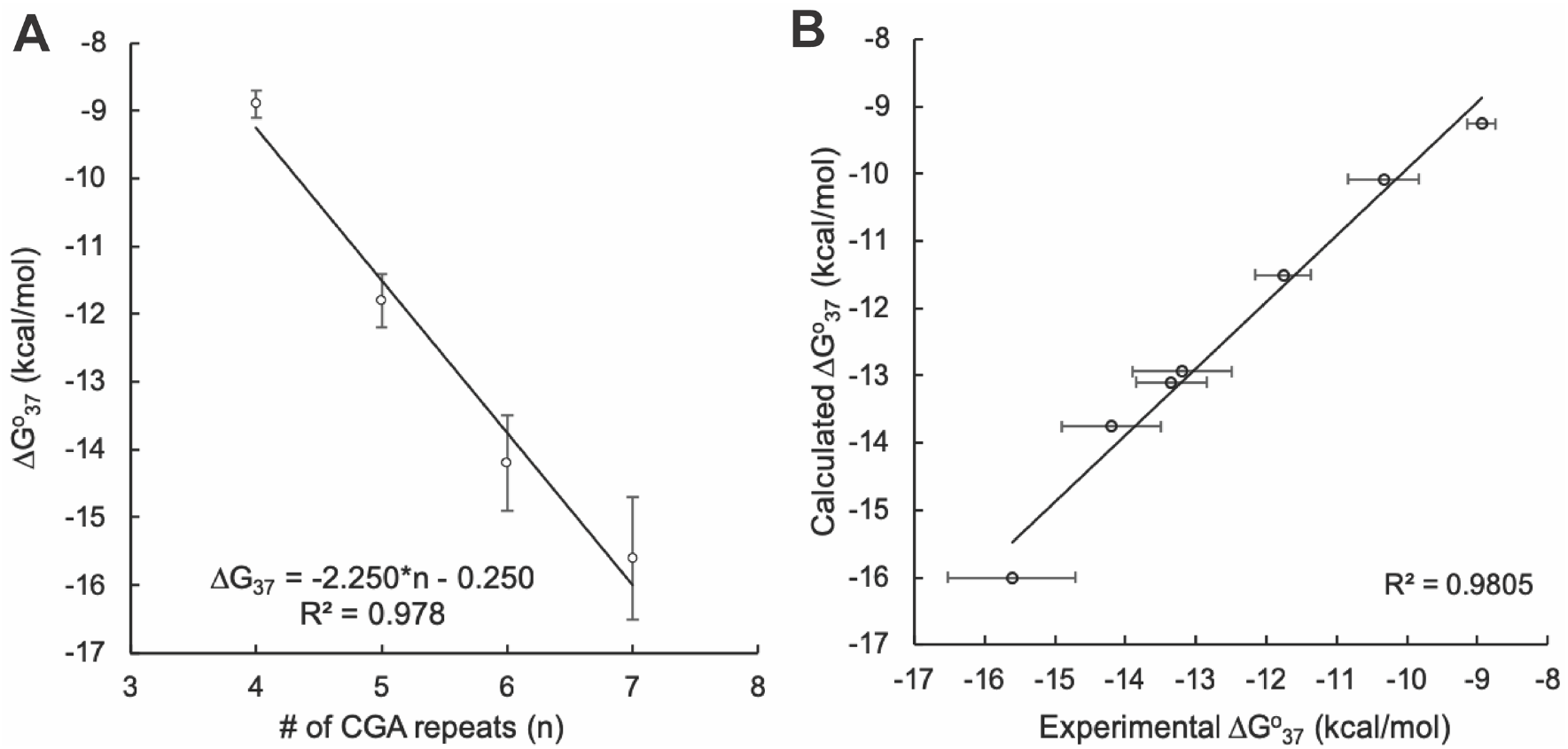
Thermodynamic estimates can be used to predict ΔG°_37_ values for new triplet repeat sequences. (A) Linear fit of (CGA)_n_ sequences vs experimentally obtained ΔG°_37_ values. (B) Experimental ΔG°_37_ is highly correlated to calculated ΔG°_37_ for all sequences, assuming additive contributions from each triplet. Estimates for d(CGA) were obtained from the fit in A, while estimates for 5′-d(GGA), 5′-d(TGA), 3′-d(TGA) were obtained from Table 1. All sequences were included except (TGA)_6_ and (CGATGA)_3_.

**FIGURE S6.**
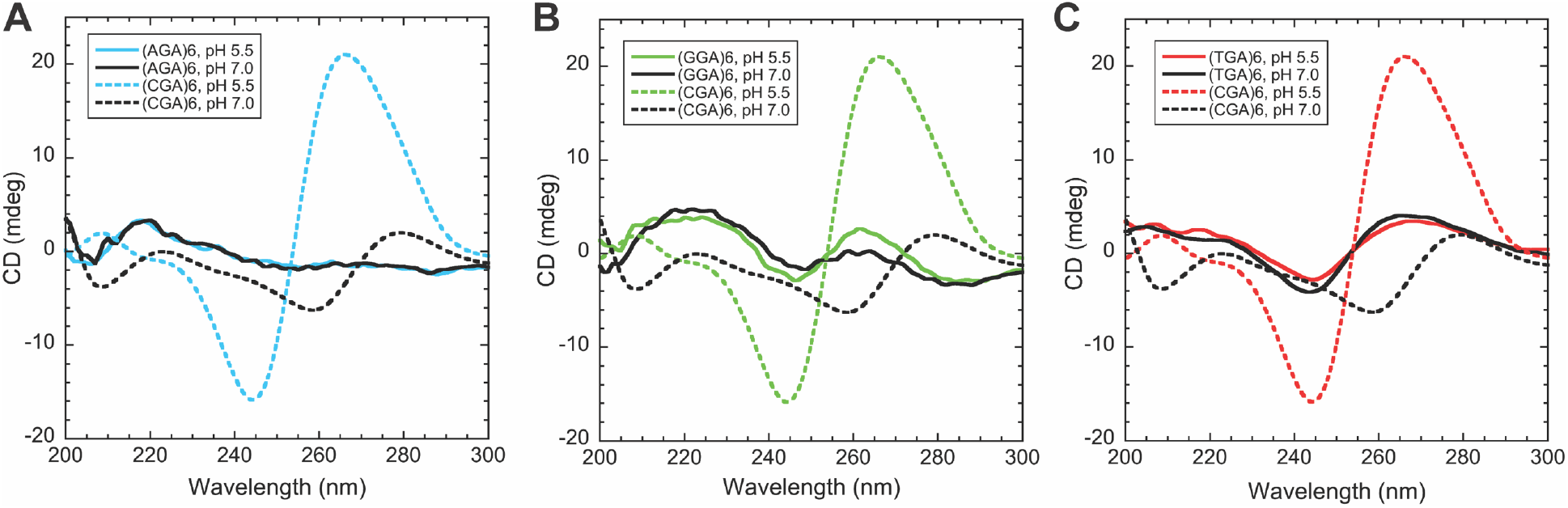
CD spectra of 10 μM (NGA)_6_ sequences compared to (CGA)_6_. (A) (AGA)_6_ does not have CD bands characteristic of parallel- or anti-parallel oriented strands and is not pH dependent. The lack of CD signal could suggest a lack of structure at each pH. (B) (GGA)_6_ also does not have CD bands characteristic of parallel- or anti-parallel oriented strands and is not pH dependent. The band at 220 nm could indicate the formation of a different structure at each pH. (C) (TGA)_6_ has CD signal characteristic of the anti-parallel form at each pH, suggesting a lack of pH sensitivity.

**FIGURE S7.**
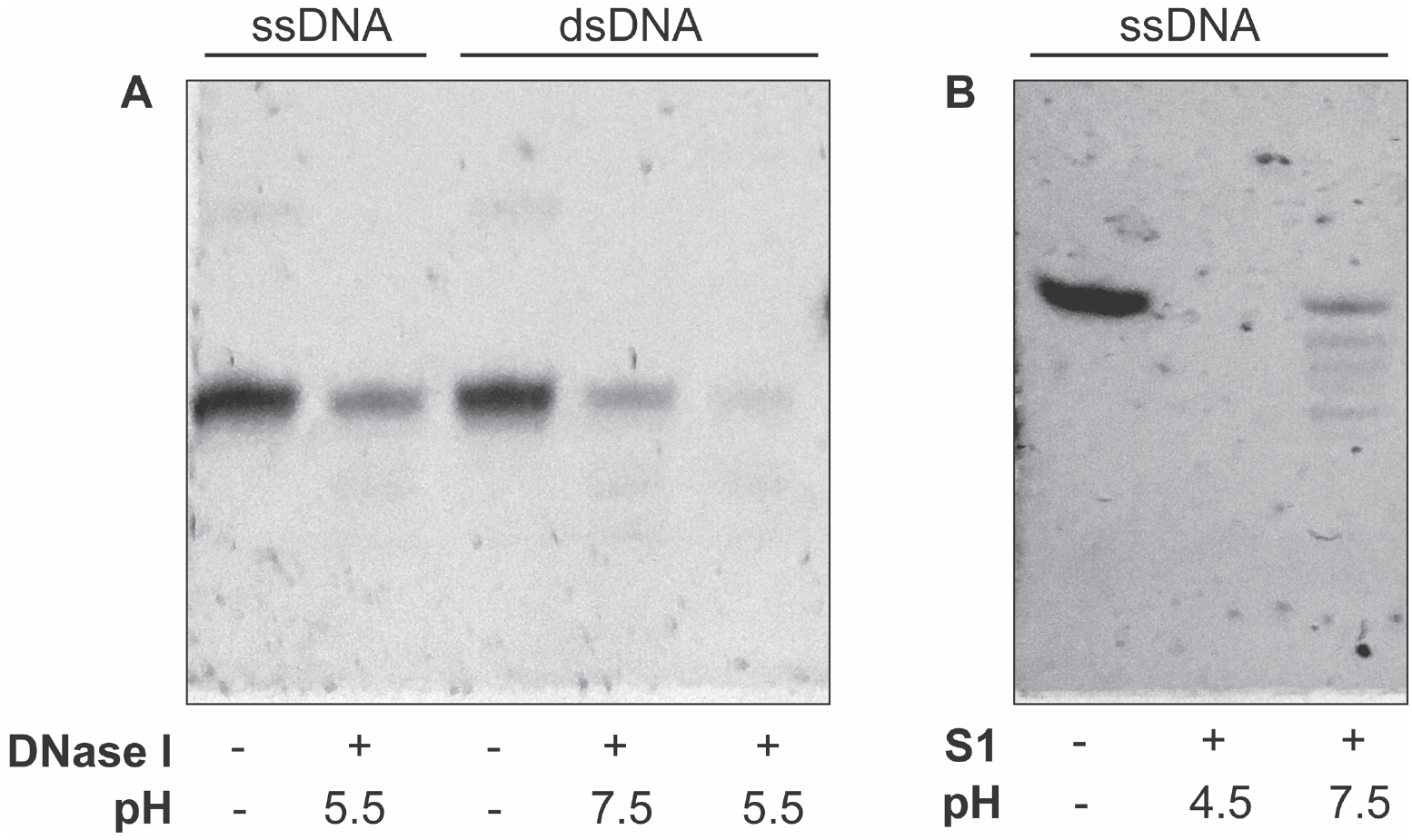
Control DNase I and S1 nuclease sensitivity experiments performed on d(GGACAGCTGGGAG) with Mg^2+^ to form double stranded (ds) DNA or without Mg^2+^ for single stranded (ss) DNA. (A) ssDNA and dsDNA exposed to DNase I at pH 5.5 and 7.5. DNase I functioned optimally on dsDNA at pH 5.5. Reduced activity was observed for ssDNA at pH 5.5 and dsDNA at pH 7.5. (B) Control ssDNA incubated with S1 nuclease at pH 4.5 and 7.5. S1 nuclease was more active at pH 4.5 than at pH 7.5.

**FIGURE S8.**
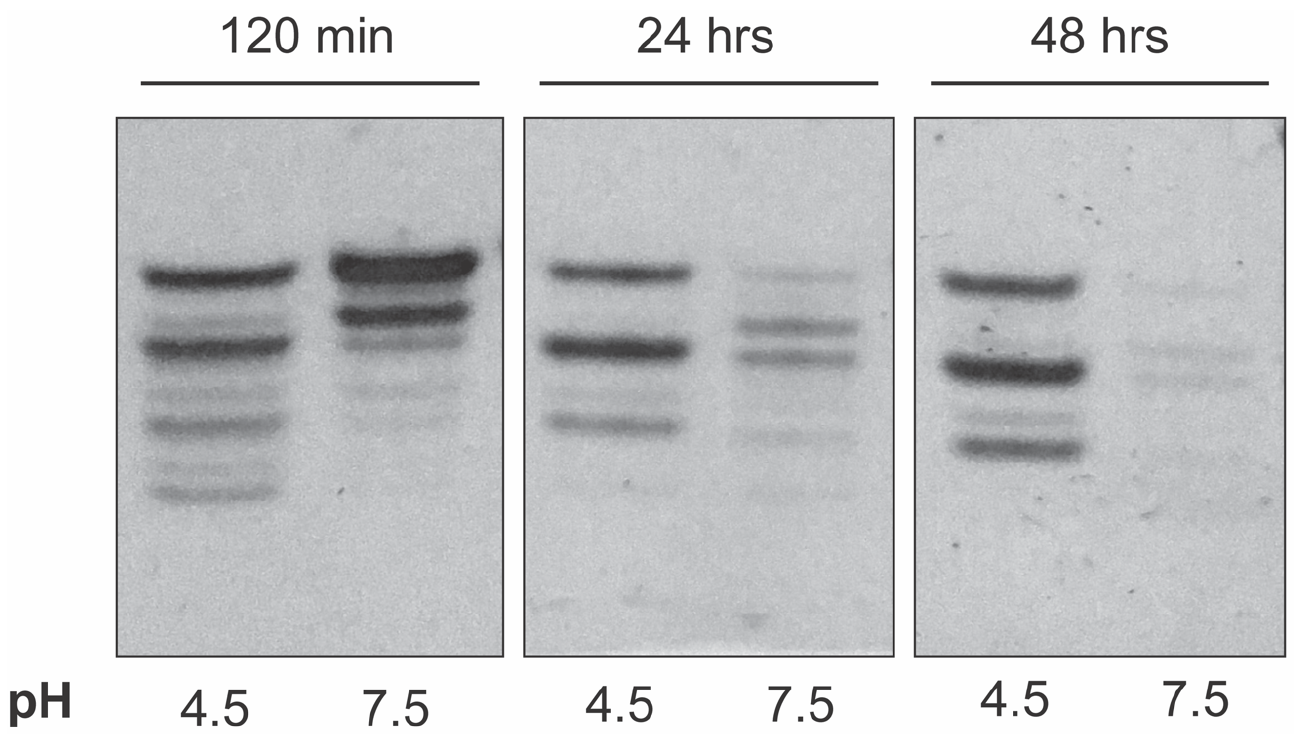
Extended S1 digestion of TGA(CGA)_5_. The bands are almost completely digested at pH 7.5 when TGA(CGA)_5_ was exposed to S1 nuclease for 2 days, suggesting that the structure formed at this pH is dynamic or not resistant to S1 digestion over time. In contrast, the intensity of the 18, 15, and 12-nt bands corresponding to digestion by S1 nuclease at pH 4.5 does not change over two days of exposure. This suggests that the parallel-stranded duplex formed is stable and resistant to S1 degradation at room temperature.

**FIGURE S9.**
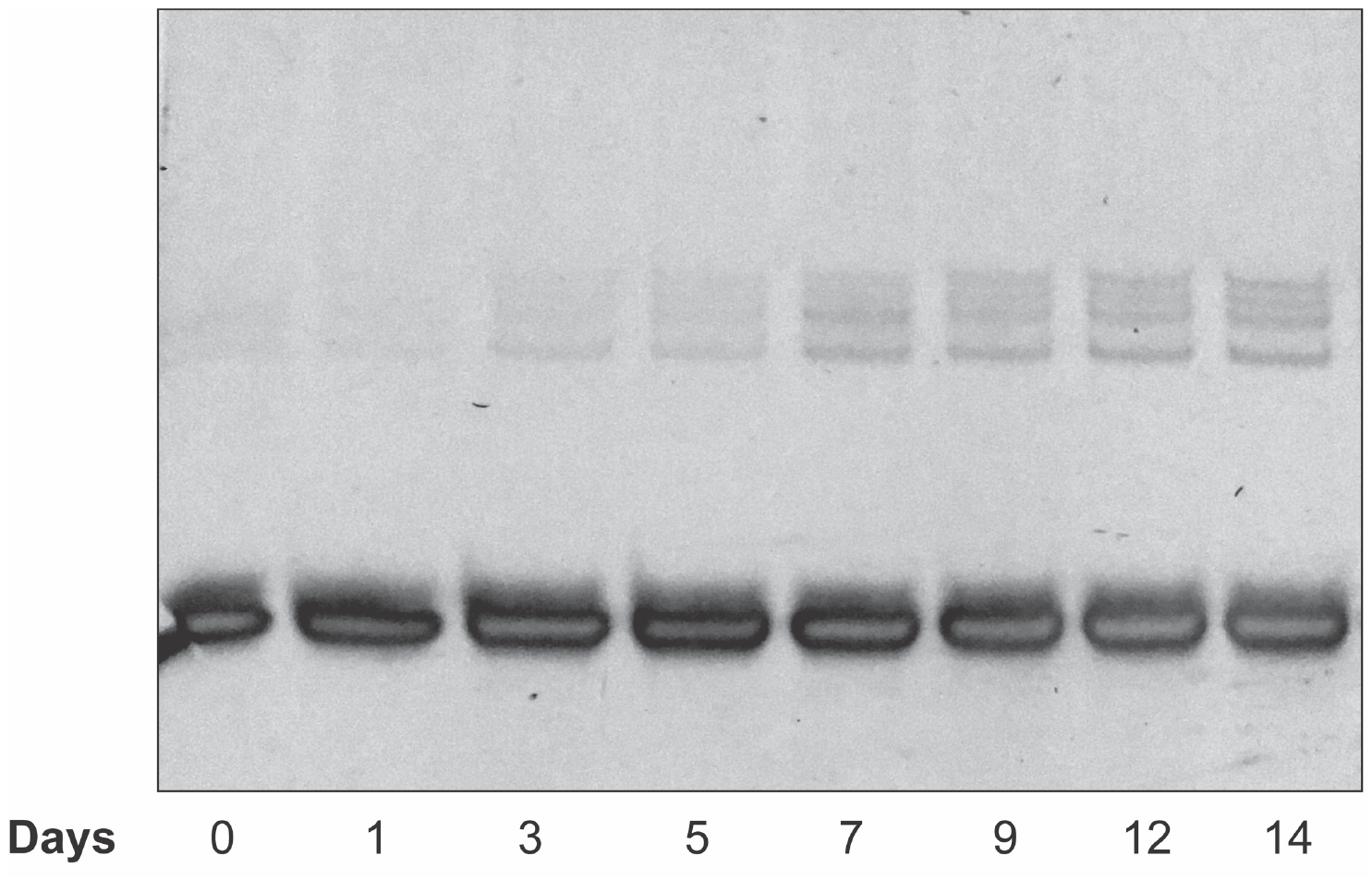
High molecular weight (HMW) bands formed over time when (CGA)_6_ is incubated at room temperature for up to 2 weeks, as shown by denaturing PAGE. All variants, except (TGA)_6_, contain HMW bands in 8 M urea denaturing gels (not shown). HMW band formation increases with time and storage temperature.

